# Independent Yet Synergistic Roles of Synaptotagmin-1 and Complexin in Calcium Regulated Neuronal Exocytosis

**DOI:** 10.1101/2019.12.16.878686

**Authors:** Sathish Ramakrishnan, Manindra Bera, Jeff Coleman, James E. Rothman, Shyam S. Krishnakumar

## Abstract

Calcium (Ca^2+^)-evoked release of neurotransmitters from synaptic vesicles requires mechanisms both to prevent un-initiated fusion of vesicles (clamping) and to trigger fusion following Ca^2+^-influx. The principal components involved, namely the vesicular fusion machinery (SNARE proteins) and the regulatory proteins (Synaptotagmin-1 and Complexin) are well-known. Here, we use a reconstituted single-vesicle fusion assay to delineate a novel mechanism by which Synaptotagmin-1 and Complexin act independently but synergistically to establish Ca^2+^-regulated fusion. Under physiologically-relevant conditions, we find that Synaptotagmin-1 oligomers bind and clamp a limited number of ‘central’ SNARE complexes via the primary binding interface, to introduce a kinetic delay in vesicle fusion mediated by the excess of free SNAREpins. This in turn enables Complexin to independently arrest the remaining free ‘peripheral’ SNAREpins to produce stably clamped vesicles. Activation of the central SNAREpins associated with Synaptotagmin-1 by Ca^2+^ is sufficient to trigger rapid (<100 msec) and synchronous fusion of the docked vesicles.

## INTRODUCTION

The controlled yet rapid (sub-millisecond) release of neurotransmitters stored in synaptic vesicles (SV) is central to all information processing in the brain (Sudhof 2013; Kaeser and Regehr 2014; Rizo 2018). Synaptic release of neurotransmitters relies on the efficient coupling of SV fusion to the triggering signal – action potential (AP)-evoked Ca^2+^ influx into the pre-synaptic terminal (Kaeser and Regehr 2014; Sudhof 2013). SV fusion is catalyzed by synaptic SNARE (soluble N-ethylmaleimide-sensitive factor attachment protein receptor) proteins, VAMP2 on the vesicles (v-SNAREs) and Syntaxin/SNAP25 (t-SNAREs) on the pre-synaptic membrane (Sollner et al. 1993; Weber et al. 1998). On their own SNARE proteins are constitutively active and intrinsically trigger fusion in the range of 0.5-1 sec (Ramakrishnan et al. 2018; Ramakrishnan et al. 2019; Xu et al. 2016).

To achieve the requisite speed and precision of synaptic transmission, the nerve terminals maintain a pool of docked vesicles that can be readily released upon Ca^2+^ influx (Sudhof 2013; Kaeser and Regehr 2014; Sudhof and Rothman 2009). The prevailing theory is that in a ‘release-ready’ vesicle, multiple SNARE complexes are firmly held (“clamped”) in a partially assembled state. These ‘SNAREpins’ are then synchronously released by Ca^2+^ to drive fusion dramatically faster than any one SNARE alone (Rothman et al. 2017; Brunger et al. 2018; Volynski and Krishnakumar 2018). Several lines of evidence suggest that two synaptic proteins, Synaptotagmin-1 (Syt1) and Complexin (Cpx) play a critical role in establishing Ca^2+^-regulated neurotransmitter release (Geppert et al. 1994; Xu, Mashimo, and Sudhof 2007; Bacaj et al. 2013; Yang, Cao, and Sudhof 2013; Huntwork and Littleton 2007).

Synaptotagmin-1 is a SV-localized protein with a large cytoplasmic part containing tandem C2A and C2B domains that bind Ca^2+^ (Fuson et al. 2007; Sutton et al. 1995) It is well established that fast AP-evoked synchronous release is triggered by Ca^2+^ binding to Syt1 C2 domains (Brose et al. 1992; Geppert et al. 1994; Littleton et al. 1993). Genetic analysis shows that Syt1 also plays a crucial role in ‘clamping’ spontaneous release and Ca^2+^-evoked asynchronous release to ensure high fidelity of the Ca^2+^-coupled synchronous transmitter release (Xu et al. 2009; Bacaj et al. 2013; Littleton et al. 1993). Recently it has been demonstrated that C2B-driven self-oligomerization of Syt1 provides the structural basis for its clamping function (Wang et al. 2014; Bello et al. 2018; Tagliatti et al. 2019). Syt1 is also involved in initial stage of docking of SVs to the plasma membrane (PM), in part mediated by its interaction with the anionic lipids, phosphatidylserine (PS) and phosphatidylinositol 4, 5-bisphosphate (PIP2) on the plasma membrane (PM) (Honigmann et al. 2013; Parisotto et al. 2012; Park et al. 2012). Besides the membrane interaction, the Syt1 also binds the neuronal t-SNAREs on the PM both independently (primary binding site) and in conjunction with Complexin (tripartite binding site) (Zhou et al. 2015; Zhou et al. 2017; Grushin et al. 2019). The Syt1-SNARE interactions are important for Syt1 role in SV docking, clamping fusion and triggering Ca^2+^-dependent neurotransmitter release (Zhou et al. 2015; Zhou et al. 2017)

Complexin is a cytosolic α-helical protein that binds and regulates SNAREpins assembly (Chen et al. 2002; Kummel et al. 2011; Li et al. 2011). Biochemical analyses show that Cpx catalyzes the initial stages of SNARE assembly, but then blocks complete assembly (Giraudo et al. 2009; Kummel et al. 2011; Li et al. 2011). Physiological studies show that Cpx indeed clamps spontaneous fusion in invertebrate neurons (Huntwork and Littleton 2007; Wragg et al. 2013; Cho et al. 2014), but its importance as a fusion clamp in mammalian neurons is still under debate (Yang, Cao, and Sudhof 2013; Lopez-Murcia et al. 2019; Courtney et al. 2019). However, under all conditions, Cpx has been shown to facilitate vesicle priming and promote AP-evoked synchronous release (Yang, Cao, and Sudhof 2013; Yang et al. 2015; Lopez-Murcia et al. 2019).

It is broadly accepted that Syt1 and Cpx are both involved in clamping spurious or delayed fusion events and synchronize neurotransmitter release to Ca^2+^-influx. However, there is presently no coherent understanding how these proteins assemble and operate together. To gain mechanistic insights into this process, a reductionist approach where the variables are limited, and components can be rigorously controlled is required. We recently described a biochemically-defined fusion system based on a pore-spanning lipid bilayer setup that is ideally suited for this purpose (Ramakrishnan et al. 2018). This reconstituted setup is capable of precisely tracking individual vesicle docking, clamping (delay from docking to spontaneous fusion) and Ca^2+^ triggered release at tens of milliseconds timescale (Coleman et al. 2018; Ramakrishnan et al. 2019). Critically, this setup allows us to examine these discrete sub-steps in the vesicular exocytosis process, independent of alterations in the preceding or following stages (Coleman et al. 2018; Ramakrishnan et al. 2019; Ramakrishnan et al. 2018).

Using the *in vitro* fusion system, we recently reported that under artificially low-VAMP2 conditions (∼13 copies of VAMP2 and ∼25 copies of Syt1 per vesicle), Syt1 oligomerization is both necessary and sufficient to establish a Ca^2+^-sensitive fusion clamp (Ramakrishnan et al. 2019). Here we extend this study to more physiologically-relevant conditions, using synaptic vesicle mimics reconstituted with ∼25 copies of Syt1 and ∼70 copies of VAMP2. We report that under these conditions, independent action of both Cpx and Syt1 oligomers are needed to stably clamp all SNARE complexes and the reversal of the Syt1 clamp is sufficient to achieve fast, Ca^2+^-triggered synchronized fusion.

## RESULTS

With the goal of approximating the physiological context, we chose reconstitution condition for small unilamellar vesicles (SUV) resulting in an average of 74 copies and 25 copies of outward-facing VAMP2 and Syt1 respectively (Figure 1 Supplement 1). We employed pre-formed t-SNAREs (1:1 complex of Syntaxin1 and SNAP-25) in the planar bilayers (containing 15% PS and 3% PIP2) to both simplify the experimental approach and to bypass the requirement of SNARE-assembling chaperones, Munc18 and Munc13 (Baker and Hughson 2016; Rizo 2018). In most of the experiments, we used fluorescently-labelled lipid (2% ATTO647-PE) included in the SUVs to track the docking, diffusion and fusion of individual SUVs (Figure 1A).

**Figure 1.**
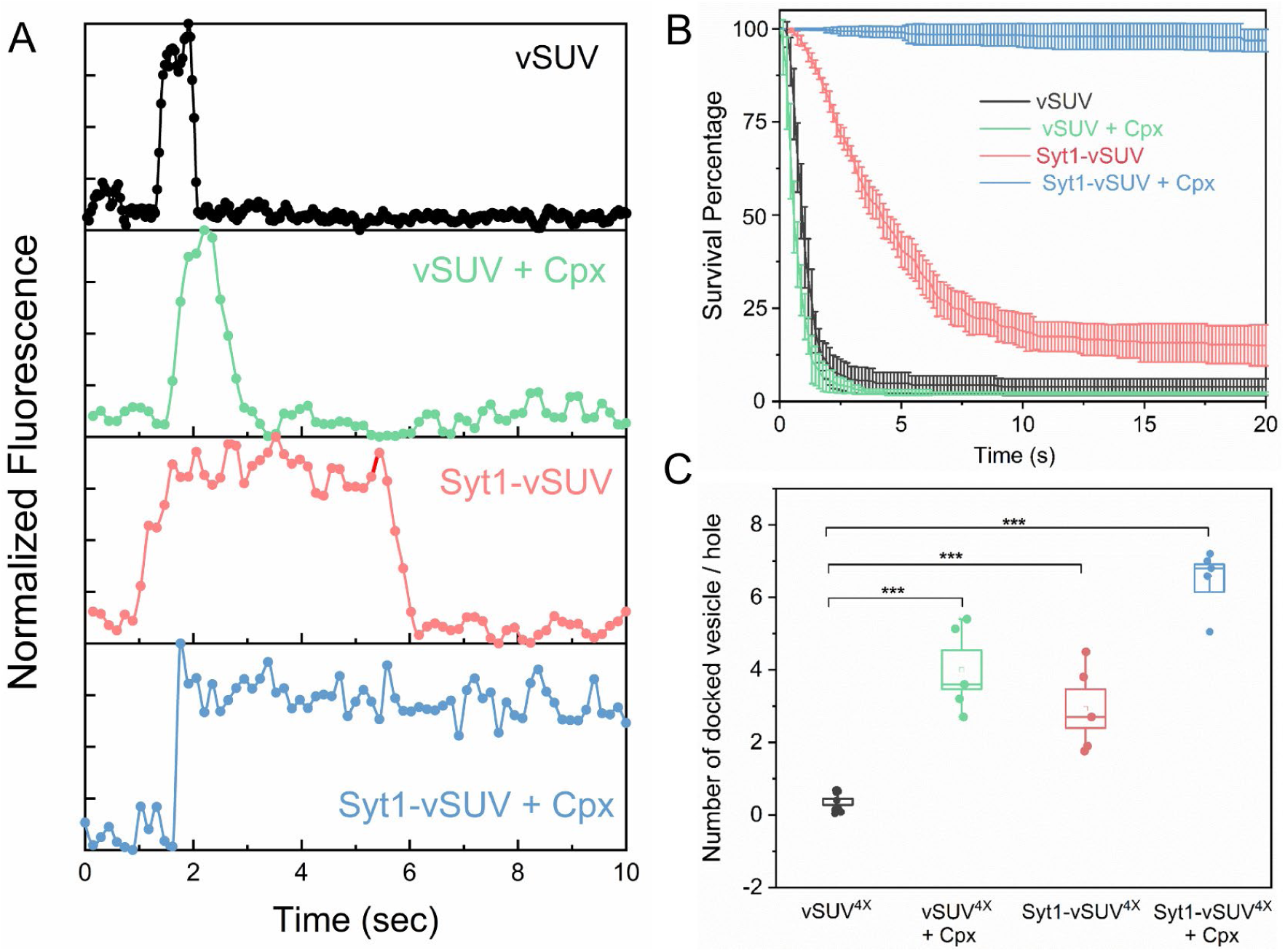
Syt1 and Cpx co-operatively clamp SNARE-mediated vesicle fusion under physiologically-relevant conditions. The effect of Syt1 and Cpx on SNARE-driven fusion was assessed using a single-vesicle docking and fusion analysis with a pore-spanning bilayer setup (Ramakrishnan et al. 2019; Ramakrishnan et al. 2018). (A) Representative fluorescence (ATTO647N-PE) traces showing the behavior of small unilamellar vesicles (SUV) containing VAMP2 (vSUV) or Syt1 and VAMP2 (Syt1-vSUV) on t-SNARE containing bilayer in the presence or the absence of Cpx. (B) The time between docking and fusion was measured for each docked vesicle and the results for the whole population are presented as a survival curve. vSUVs (black curve) are diffusively mobile upon docking (Figure 1 Supplement 2) and fuse spontaneous with a half-time of ∼1 sec. Addition of soluble Cpx (2 μM) does not change this behavior (green curve). Inclusion of Syt1 in the v-SUV (red curve) does not block fusion but increases the time from docking-to-fusion (∼ 5 sec half-life), in effect delaying the kinetics of fusion. When included together Syt1 and Cpx (blue curves) fully arrest fusion to produce stably docked SUVs that attach and remain in place during the entire period of observation. (C) Syt1 and Cpx, both individually and collectively increase the number of docked vesicles. In all cases, a mutant form of VAMP2 (VAMP2^4X^) which eliminated fusion was used to unambiguously estimate the number of docked vesicles after the 10 min interaction phase. The average values and standard deviations from 3-4 independent experiments are shown for each condition. In sum, 500-1000 vesicles were analyzed for each condition.

We initially focused on the kinetics of constitutive fusion to assess the ability of Syt1 and Cpx to ‘clamp’ SNARE-driven fusion in the absence of Ca^2+^. We monitored large ensembles of vesicles to determine the percent remaining unfused as a function of time elapsed after docking and quantified as ‘survival percentages’ (Figure 1B). Docked immobile vesicles that remained un-fused during the initial 10 min observation period were defined as ‘clamped’ (Ramakrishnan et al. 2019). Vesicles containing VAMP2 only (vSUV) that docked to the t-SNARE containing bilayer surface were mobile and a majority (>95%) spontaneously fused typically with a t_1/2_ ∼ 1 sec post-docking (Figure 1B, Figure 1 Supplement 2).

Inclusion of wild-type Syt1 in the vesicles (Syt1-vSUVs) enhanced the vesicle docking rate, with ∼8-fold increase in total number of docked vesicles (Figure 1C). The majority (∼80%) of docked Syt1-vSUVs remained mobile on the bilayer surface and fused on an average ∼5-6 sec after docking (Figure 1B, Figure 1 Supplement 2, Video 1). The remainder small fraction (∼20%) were immobile and stably clamped. This is in stark contrast to our earlier finding under low-copy VAMP2 conditions wherein the bulk of the Syt1-vSUVs (>90%) were stably clamped (Ramakrishnan et al. 2019). Nonetheless, the pronounced docking-to-fusion delay introduced by Syt1 (t_1/2_ ∼5 sec for Syt1-vSUV *compared to* ∼1 sec for vSUV) suggests that under physiologically-relevant (“normal” VAMP copy number) conditions, Syt1 alone can meaningfully delay but not stably clamp fusion.

This unstable Syt1 clamp was stabilized by addition of Cpx (Figure 1B, Figure 1 Supplement 2, Video 2). In the presence of 2 μM of soluble Cpx, all Syt1-vSUVs were immobile following docking (Figure 1B), and they rarely fused over the initial observation period (Figure 1B, Figure 1 Supplement 2, Video 2). In fact, these vesicles remained stably docked up to 45 min without fusing (Figure 1 Supplement 3). Furthermore, Syt1 and Cpx together significantly increased (∼18-fold) the total number of docked vesicles (Figure 1C). Thus, we find that Syt1 and Cpx act synergistically to increase the rate of vesicle docking but block fusion of the docked vesicles under resting conditions.

Addition of soluble Cpx (2 μM) alone produced a ∼10-fold increase in the number of docked vesicles (Figure 1C) but did not change the behavior of the docked vSUVs (Figure 1B, Figure 1 Supplement 2). In the presence of Cpx alone, virtually all docked vSUVs fused spontaneously typically within 1-2 sec (Figure 1A, B). This meant that Syt1 is somehow required for Cpx to function as a fusion clamp.

We then investigated the effect of Ca^2+^ on the stably Syt1/Cpx-clamped vesicles (Figure 2). We estimated the time of arrival of Ca^2+^ at/near the docked vesicles using a lipid-conjugated Ca^2+^ indicator (Calcium green C24) attached to the planar bilayer (Figure 2 Supplement 1). Influx of free Ca^2+^ (1 mM) triggered simultaneous fusion of >90% of the docked vesicles (Figure 2B, Video 3). These vesicles fused rapidly and synchronously, with a characteristic time-constant (τ) of 0.11 sec following the arrival of Ca^2+^ locally (Figure 2C). This estimate is constrained by the temporal resolution limit (150 msec per frame) of our imaging experiment. Indeed, most of Ca^2+^-triggered fusion occurs within a single frame (Video 3). We thus believe that the true Ca^2+^-driven fusion rate is likely <100 msec. Our data indicates that Syt1 and Cpx acting together synchronize vesicle fusion to Ca^2+^ influx and greatly accelerates the underlying SNARE-mediated fusion process (which typically occurs at a rate of ∼1 sec). We also tested and confirmed these findings with a content-release assay using sulforhodamine B-loaded SUVs (Figure 2 Supplement 2) under similar experimental conditions. Overall, we find that Syt1 and Cpx act co-operatively to clamp the SNARE assembly process to generate and maintain a pool of docked vesicles that can be triggered to fuse rapidly and synchronously to Ca^2+^ influx. In other words, Syt1 and Cpx are necessary and sufficient to establish Ca^2+^-regulated exocytosis.

**Figure 2.**
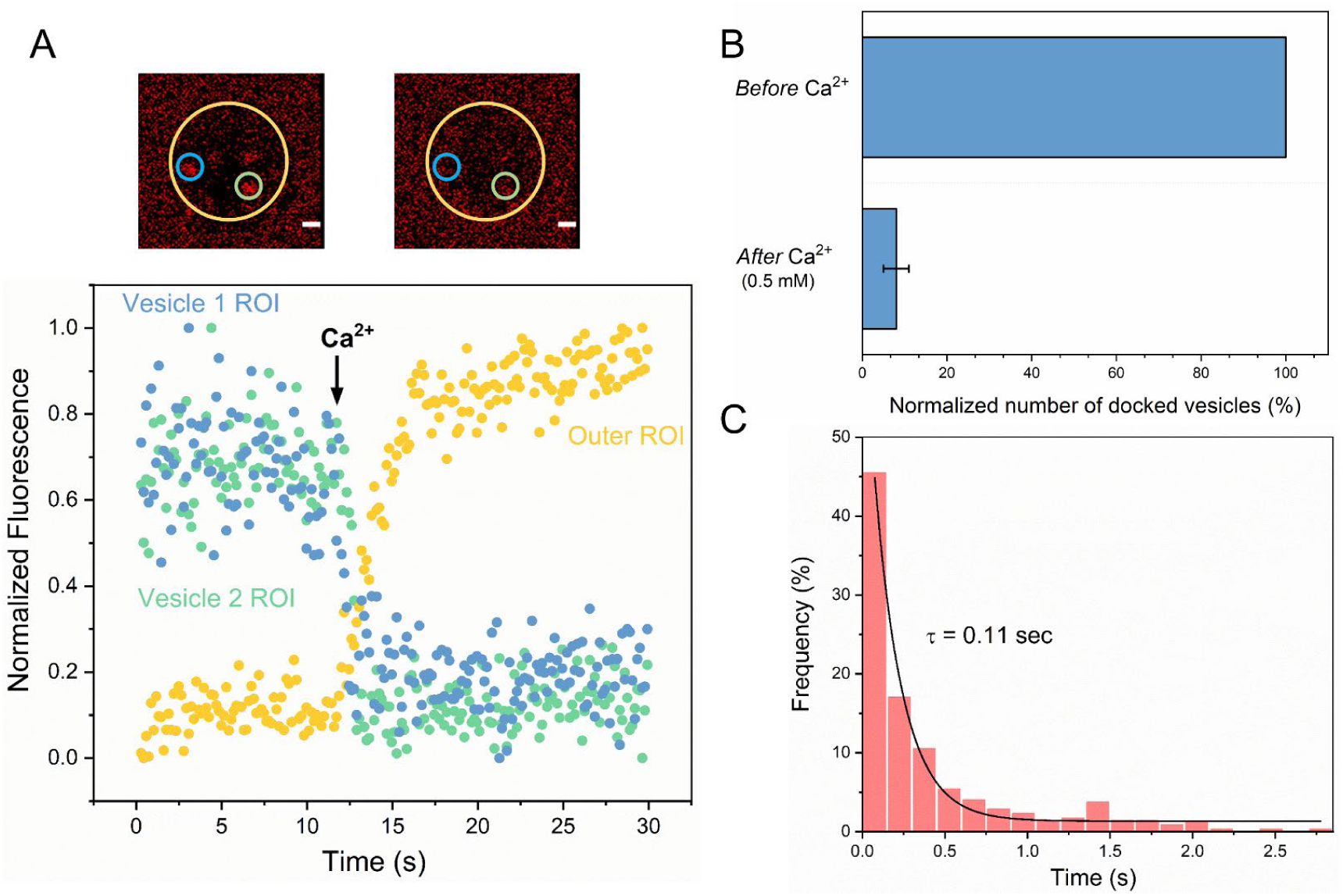
Syt1/Cpx clamped vesicles fuse synchronously and rapidly following Ca^2+^-addition. (A) Representative fluorescence images (top) and quantitation of change in fluorescence signal (bottom) before and after addition of 1 mM Ca^2+^ shows that vesicles clamped by Syt1/Cpx are sensitive to Ca^2+^. Fusion was attested by a burst and sudden decrease in fluorescence (ATTO647N-PE) intensity as the lipids diffuse away. To visualize this, fluorescence was simultaneously monitored in a circular region of interest (ROI) encompassing the docked vesicle (vesicle ROI, green and blue circles) and in a surrounding annular ROI (outer ROI, yellow circle). Corresponding to actual fusion events, we observed a sudden decrease of fluorescence intensity in the vesicle ROI with a concomitant increase of fluorescence in the annular outer ROI. Note that the two docked vesicles fuse synchronously in response to Ca^2+^-influx. (B) End-point analysis at 1 min post Ca^2+^-addition shows that >90% of all clamped vesicles fuse following Ca^2+^ addition. (C) Kinetic analysis shows that the Syt1/Cpx clamped vesicles fuse rapidly following Ca^2+^-addition with a characteristic time constant of 0.11 sec. This represents the temporal resolution limit of our recordings and the true Ca^2+^-triggered fusion rate is likely well below 0.1 sec. Thus, Syt1 and Cpx synchronize vesicle fusion to Ca^2+^-influx and accelerate the underlying fusion process. The average values and standard deviations from three independent experiments (with ∼1000 vesicles in total) is shown.

We next examined if Syt1 and Cpx act on the same SNARE complex sequentially or if they function independently to produce molecularly-distinct clamped SNAREpins under the same docked vesicles. To this end, we washed Cpx out from stably clamped Syt1/Cpx vesicles (by dilution) in the absence or the presence of excess inhibitory soluble cytoplasmic domain of the t-SNARE complex (CDT). Cpx binds to half-zippered (clamped) SNAREpins with a K_d_ ∼0.5 μM (Krishnakumar et al. 2011; Kummel et al. 2011) and is therefore expected to freely dissociate when the bulk concentration of CPX is reduced well below that. CDT will bind and sequester/inactivate any free VAMP2 on the vesicles and is also expected to effectively compete against bilayer-anchored t-SNAREs. This is because CDT could form fully-zippered SNARE complexes (stabilized by ∼70 k_B_T) as compared to the half-zippered SNAREpins (∼35 k_B_T) formed by bilayer-anchored t-SNAREs (Li et al. 2016). We reasoned that If Syt1 and Cpx act on the same SNARE complex, then CDT treatment (with Cpx wash-out) would irreversibly block vesicle fusion. However, if Syt1 and Cpx clamp different SNARE complexes, then some SNAREpins might be protected from CDT and thus, the vesicles will remain sensitive to Ca^2+^ influx.

We used fluorescently-labeled Cpx to test and confirm the near-complete washout of Cpx from the clamped Syt1/Cpx vesicles following the extensive (40X) buffer wash (Figure 3 Supplement 1). Without CDT, the docked vesicles proceeded to fuse spontaneously following the buffer wash (Figure 3A, Figure 3 Supplement 2). This further confirmed that both Syt1 and Cpx are needed to produce a stably clamped state. In the presence of CDT, most of the vesicles remained docked even after the removal of previously-bound Cpx by the buffer wash (Figure 3B, Figure 3 Supplement 2). Subsequent addition of Ca^2+^ (1 mM) triggered rapid and synchronous fusion of the docked vesicles (Figure 3B, Figure 3 Supplement 2), with fusion kinetics similar to the control experiments (Figure 2C). This implied that there are at least two types of clamped SNAREpins under a docked vesicle – those clamped by Syt1 (which are shielded from CDT) and others arrested by Cpx (which become accessible to CDT following the buffer wash-out). It further indicated that even though both Syt1 and Cpx clamps are required to produce a stably ‘clamped’ vesicle, the activation of the Syt1-associated SNAREpins is sufficient to elicit rapid, Ca^2+^-synchronized vesicular fusion.

**Figure 3.**
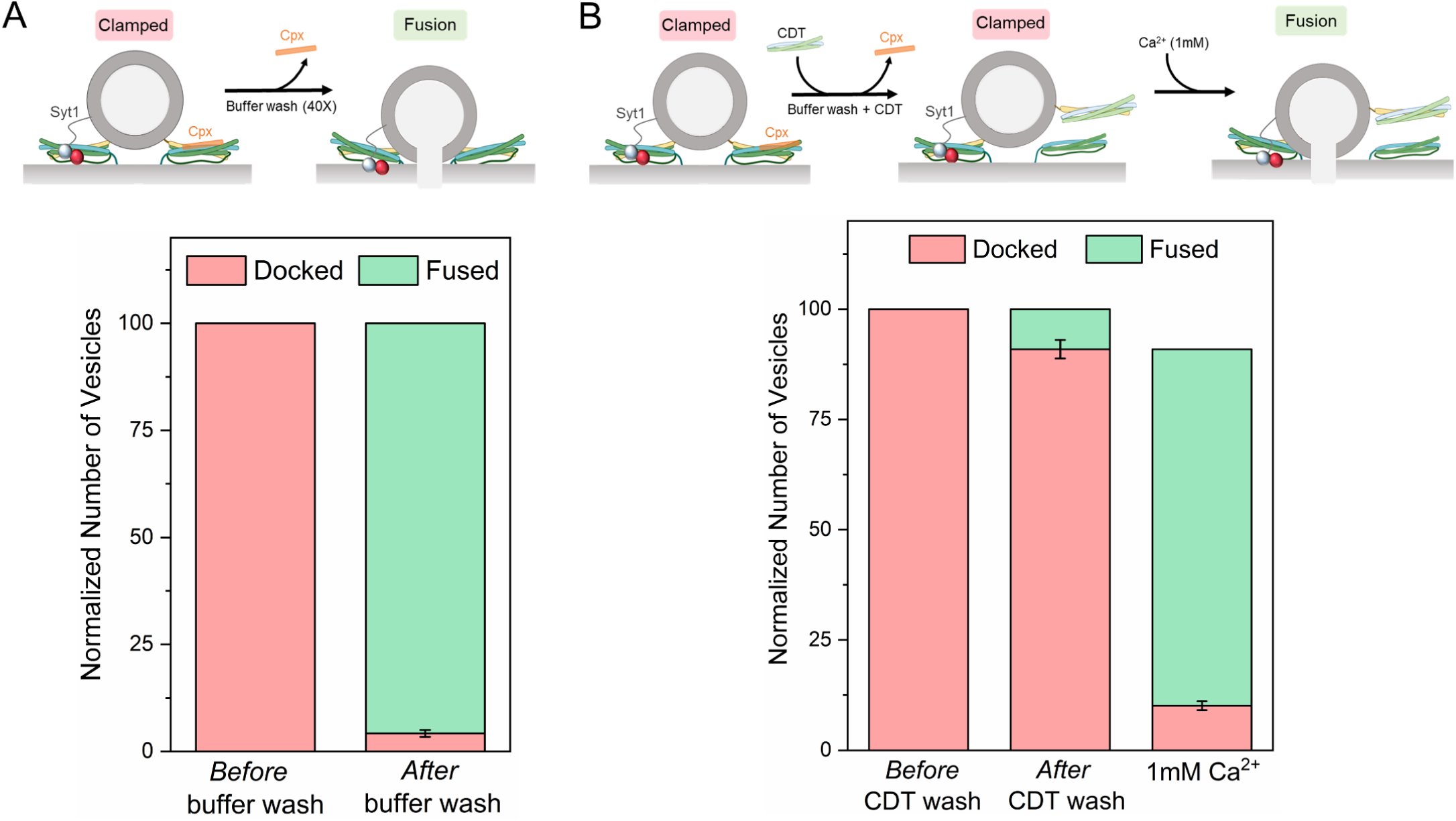
Syt1 and Cpx bind and clamp different pools of SNAREpins. (A) *In situ* removal of Cpx from the Syt1/Cpx clamped state by extensive (40X) buffer wash triggers spontaneous fusion of the docked vesicles. This further confirms that both Syt1 and Cpx are required to produce a stable clamped state. (B) Inclusion of soluble cytoplasmic domain of t-SNAREs (CDT) blocked the spontaneous fusion events triggered by elimination of Cpx from the Syt1/Cpx clamped vesicles. The CDT-treated vesicles remain sensitive to Ca^2+^ influx and most of the vesicles fuse rapidly and synchronously following the addition of 1 mM Ca^2+^. This indicates that a sub-set of SNAREpins are protected against CDT even in the absence of Cpx, implying that Syt1 and Cpx likely engage and clamp independent set of SNAREpins. It further shows that the Syt1-associated SNAREpins are sufficient to catalyze rapid Ca^2+^-triggered vesicle fusion. Data obtained from four to five independent experiments with at least 200 vesicles in total is shown. Note: Only single SNAREpins clamped by either Syt1 or Cpx is shown for illustrative purposes.

Considering these findings, we sought to establish the molecular determinants of the Syt1 clamp and its reversal by Ca^2+^. To focus on the Syt1 component of the clamp, we tested the effect of specific Syt1 mutations using low copy VAMP2 conditions, i.e. SUVs containing ∼13 copies of VAMP2 and ∼25 copies of Syt1 (wild type or mutants) in the absence of Cpx. Consistent with our earlier report (Ramakrishnan et al. 2019), wild-type Syt1 (Syt1^WT^) alone was sufficient to produce stably-clamped vesicles under these conditions (Figure 4A). Selective disruption of Syt1-SNARE ‘primary’ binding using the previously described (Zhou et al. 2015; Zhou et al. 2017) mutations in Syt1 C2B (R281A/E295A/Y338W/R398A/R399A; Syt1^Q^) and SNAP25 (K40A/D51A/E52A/E55A/D166A, SNARE^Q^) abolished the Syt1 clamp (Figure 4A), with >99% of the docked Syt1^Q^ vesicles fusing constitutively in the 10 min observation period. Destabilization of the Syt1 C2B oligomers with a point-mutation (F349A)(Bello et al. 2018) produced a similar phenotype, wherein ∼85% of docked Syt1^F349A^ vesicles proceeded to fuse spontaneously (Figure 4A). On the other hand, disrupting Ca^2+^ binding to the C2B domain (D309A, D363A, D365A; Syt1^DA^)(Shao et al. 1996) had no effect on Syt1 clamping function, with all vesicles remaining un-fused (Figure 4A and 4B). This meant that Syt1 ability to oligomerize and bind SNAREpins via the primary binding site is key to its clamping function.

**Figure 4.**
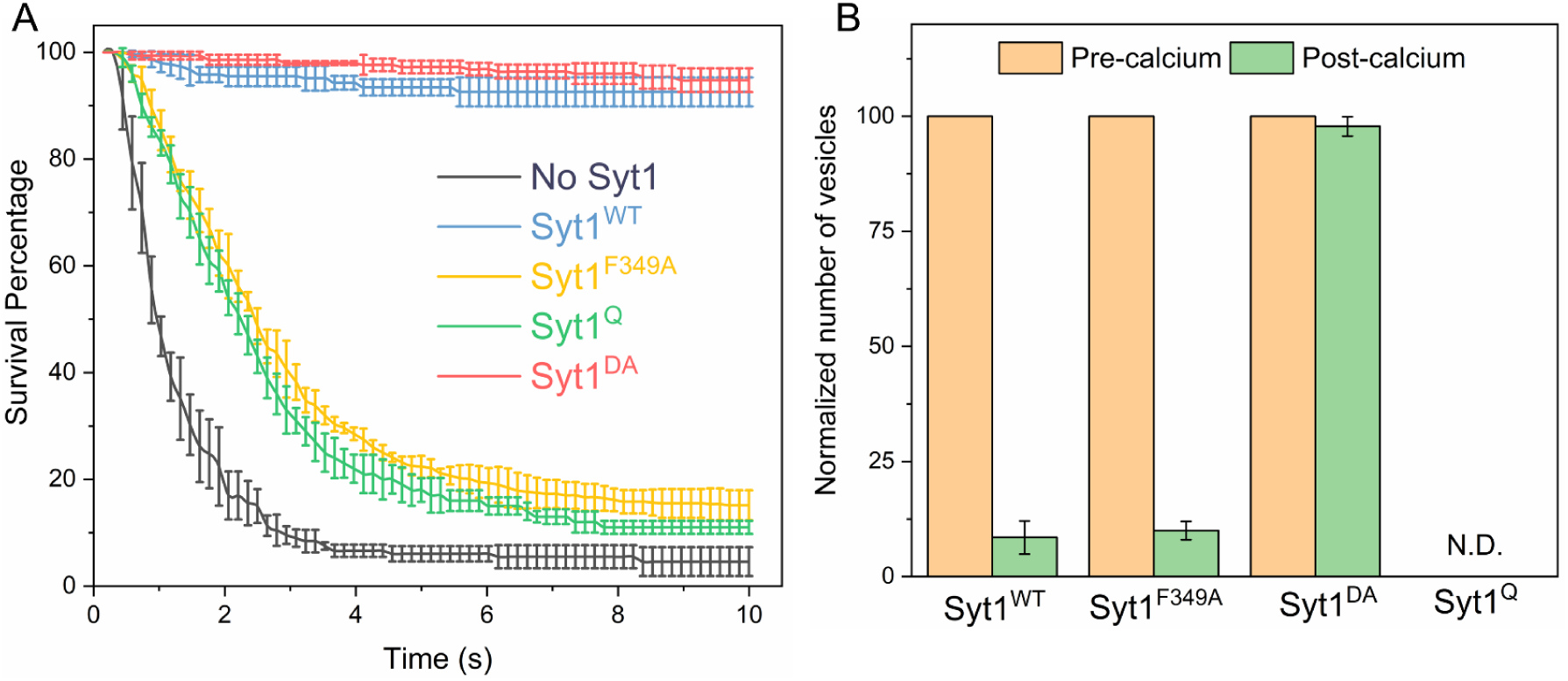
Molecular determinants of Syt1 clamp and its reversal by Ca^2+^. To focus on the Syt1 component of the fusion clamp, all fusion analysis was carried out using vesicles containing low copy VAMP2 (∼13 copies) with normal number (∼25 copies) of Syt1 molecules (wild-type or targeted mutations) in the absence of Cpx. (A) Survival analysis shows that disrupting the Syt1-SNARE primary interface (Syt1^Q^, green curve) or destabilizing Syt1 oligomerization (Syt1^F349A^, yellow curve) abrogates the Syt1 clamp, whilst Ca^2+^-binding motif on the Syt1 C2B domain (Syt1^DA^, red curve) is not involved in establishing the fusion clamp. (B) Addition of Ca^2+^ triggered rapid fusion of vast majority (>90%) of remainder of the docked Syt1^F349A^ vesicles, very similar to the behavior of the Syt1^WT^ vesicles. Predictably, blocking Ca^2+^-binding to C2B domain rendered the vesicle Ca^2+^-insensitive, with the majority of Syt1^3A^ remaining un-fused. We did not have sufficient number of docked Syt1^Q^ vesicles to do a quantitative analysis, but qualitatively, the few that remained docked failed to fuse following Ca^2+^-addition. This implies that both C2B binding to Ca^2+^ and SNAREs are required for the Ca^2+^-activation, but the ability to form oligomers is not crucial for the Ca^2+^-triggered reversal of the Syt1 clamp. The average values and standard deviations from 4 independent experiments are shown for each condition. In total, ∼250 vesicles were analyzed for each condition.

Addition of Ca^2+^ (1mM) triggered rapid and synchronous fusion of the docked Syt1^WT^ vesicles and the remaining minority fraction (∼15%) of the ‘clamped’ Syt1^349A^ containing vesicles (Figure 4B). In fact, the docked Syt1^F349A^ vesicles were indistinguishable in their behavior from Syt1^WT^ vesicles, suggesting the Syt1 oligomerization is not critical for the Ca^2+^-activation mechanism. In contrast, docked vesicles containing the Ca^2+^-binding mutant (Syt1^DA^) never fused even after Ca^2+^-addition (Figure 4B). Similarly, Ca^2+^-influx failed to trigger the fusion of the residual (∼1%) Syt1^Q^ vesicles. However, the relatively small number of docked Syt1^Q^ prior to Ca^2+^-influx precludes any meaningful quantitative analysis. Nonetheless, our data suggests that Ca^2+^-binding to the Syt1 C2B domain and its simultaneous interaction with the t-SNARE protein is required for Ca^2+^-triggered reversal of the fusion clamp.

We also tested the effect of the Syt1 mutants using vesicles containing normal VAMP2 and Syt1 (WT and mutants) copy numbers in the presence of 2 μM Cpx. For all mutations tested, we observed immobile, stably docked vesicles (Figure 4 Supplement 1). However, these vesicles were insensitive to Ca^2+^ and did not fuse following Ca^2+^ (1 mM) addition (Figure 4 Supplement 1). This suggests that in the absence of the Syt1 clamp, Cpx irreversibly blocks vesicle fusion. Taken together, our data indicates that there are at least two classes of clamped SNAREpins under every docked vesicle – a small population that is reversibly clamped by Syt1 and the remainder that is irreversibly blocked by Cpx. In other words, we find that the Syt1-associated SNAREpins are required and likely sufficient for Ca^2+^-evoked fast vesicular release.

## DISCUSSION

Here we report that Syt1 and Cpx act concomitantly to clamp SNARE-driven constitutive fusion events (Figure 1). It emerges that Syt1 and Cpx sequentially bind and clamp different pools of SNARE complexes (Figure 3). These finding taken together with the known structural and biochemical properties of Syt1 and Cpx prompts a novel ‘synergistic clamping’ mechanism. We posit that Syt1 C2B domain binds PIP2 (via the poly-lysine motif) on the plasma membrane and assembles into ring-like oligomeric structures (Bello et al. 2018; Wang et al. 2014; Zanetti et al. 2016). The Syt1 oligomers concurrently binds the t-SNAREs via the ‘primary’ interface (Zhou et al. 2015; Zhou et al. 2017; Grushin et al. 2019). The Syt1-t-SNARE interaction, which likely precedes the engagement of the v- and the t-SNAREs, positions the Syt1 such that it sterically blocks the full assembly of the associated SNAREpins (Grushin et al. 2019). The Syt1 oligomers in addition to creating a stable steric impediment, could also radially restrain the assembling SNAREpins (Grushin et al. 2019; Rothman et al. 2017).

Physiologically, all three synaptic SNAREs (Syntaxin, SNAP-25 and VAMP2) are assembled into a SNAREpin in a concerted reaction involving chaperones Munc13 and Munc18 (Baker and Hughson 2016; Rizo 2018), the need for which is by-passed in our system by pre-assembled t-SNARE complex. We imagine that these chaperones also function in a concerted manner with Syt1 ring-like oligomers and have recently advanced a speculative model outlining how these proteins can co-operate to template SNAREpin assembly (Rothman et al. 2017).

The Syt1 ring-like oligomers can bind and clamp only a small sub-set of SNAREpins (which we refer to as ‘central SNAREpins’) as the number of available SNAREpins per synaptic vesicle far exceeds the Syt1 density (∼70 copies of VAMP2 vs. ∼20 copies of Syt1). The remaining ‘peripheral’ SNAREs would then be unrestrained and can fully zipper to catalyze vesicle fusion, though at an impeded rate (Figure 1). It is tempting to speculate that the limited set of Syt1-clamped ‘central’ SNAREpins correspond physiologically to the six symmetrically organized protein densities that underlie each docked synaptic-like vesicle visualized by cryo-electron tomography analysis (Li et al. 2019). Indeed, this symmetrical organization is templated by Syt1 ring-like oligomers (Li et al. 2019)

Cpx binds the SNARE complex ‘only’ when the Syntaxin and VAMP2 are partially-assembled (Chen et al. 2002; Kummel et al. 2011) and competitively blocks the complete zippering of the C-terminal portion of the VAMP2 SNARE motif (Kummel et al. 2011; Li et al. 2011; Giraudo et al. 2009). It is well-established that the c-terminal portion of VAMP2 assembles with extraordinarily high energy and rate acting as the major power stroke to drive membrane fusion (Gao et al. 2012). Consequently, Cpx, on its own, is ineffective in clamping SNARE-driven vesicle fusion (Figure 1).

We envision that under physiological conditions, Syt1 clamp sets the stage for Cpx to bind and clamp the ‘peripheral’ SNARE complexes. Syt1, in addition to fully-arresting the central SNAREpins, also hinders the assembly of the ectopic SNAREpins enabling Cpx to effectually arrest their complete zippering. In this manner, the progressive action of Syt1 and Cpx is involved in the generation of the stably docked, release-ready vesicles. Under each vesicle, Syt1 oligomers binds and clamps a small sub-set of available SNAREpins at the early stages of docking, which in turn enables Cpx to clamp the remainder of the SNAREpins. This molecular model could explain the indispensable role of both Syt1 and Cpx in establishing the fusion clamp as evidenced in several genetic deletion studies (Bacaj et al. 2013; Geppert et al. 1994; Cho et al. 2014; Martin et al. 2011; Littleton et al. 1993; Yang, Cao, and Sudhof 2013)

We further discovered that the reversal of the Syt1 clamp is sufficient to drive, rapid Ca^2+^-triggered fusion of the docked vesicles (Figure 2). It involves Syt1 C2B domain binding both Ca^2+^ and the SNARE complex (Figure 4). We have recently demonstrated that Ca^2+^-binding to Syt1 C2 domains induce a large-scale conformational rearrangement of the Syt1-SNARE complex on the lipid membrane surface, disrupting the pre-fusion clamped architecture (Grushin et al. 2019). This, in effect, reverses the Syt1 clamp, allowing the associated SNAREs to complete zippering and drive fusion. Our data further suggests that only a small fraction of available SNAREpins per vesicle are involved in the Ca^2+^-activation process. This is consistent with the earlier reports that 2-3 SNARE complexes are sufficient to facilitate Ca^2+^-evoked synchronous neurotransmitter release (Sinha et al. 2011; Mohrmann et al. 2010). Indeed, recent modeling studies considering the concept of mechanical coupling have predicted that an optimum of 4-6 SNAREpins is required to achieve sub-millisecond vesicular release (Manca et al. 2019). Our data suggests that Cpx irreversibly arrests SNAREpin assembly (Figure 4 Supplement 1) and raises the intriguing possibility that the Cpx clamp is not necessarily reversed during the Ca^2+^-activation process. This is however speculative and additional research is required to address this in detail.

Overall, our data is in alignment with the emergent view that Syt1 plays a pivotal role in orchestrating Ca^2+^-regulated neurotransmitter release. Syt1 functions both as a fusion clamp and the principal Ca^2+^-sensor to establish Ca^2+^-regulation of vesicular fusion (Geppert et al. 1994; Littleton et al. 1993). Furthermore, Syt1, by virtue of self-assembling into oligomeric structures, also provides the molecular framework to organize the exocytic machinery into a co-operative structure to enable ultra-fast fusion (Li et al. 2019; Rothman et al. 2017; Volynski and Krishnakumar 2018).

We have articulated the simplest hypothesis considering discrete ‘central’ and ‘‘peripheral’ SNAREpin associated with Syt1 and Cpx respectively. However, it is easy to imagine that Cpx could also bind the central SNAREpins. Indeed, recent X-ray crystal structure revealed that both Syt1 and Cpx can bind the same pre-fusion SNARE complex (Zhou et al. 2017). Importantly, Cpx binding has been shown to create a new binding interface on the SNAREpins (tripartite site), which could be potentially occupied by Syt1 or other Syt isoforms (Rothman et al. 2017; Volynski and Krishnakumar 2018). Such an arrangement would essentially augment the Syt1 clamp. It also could provide the structural basis to describe the synergistic regulation of neurotransmitter release by different Syt isoforms (Rothman et al. 2017; Volynski and Krishnakumar 2018) and further explain how Cpx could promote Ca^2+^-evoked synchronous neurotransmitter release. We thus, favor such a composite model, wherein Syt1 and Cpx co-operate to both clamp un-initiated fusion and activate different modes of Ca^2+^-evoked neurotransmitter release. It is worth noting that our current data does not preclude this possibility. However, further studies involving a more detailed mutational analysis, with high-temporal resolution is needed to establish the precise molecular organization of the fusion clamp and to discern the role of Cpx in promoting Ca^2+^-triggered release.

## MATERIALS AND METHODS

### Materials

The following cDNA constructs, which have been previously described (Krishnakumar et al. 2013; Krishnakumar et al. 2011; Weber et al. 1998; Mahal et al. 2002), were used in this study: full-length VAMP2 (VAMP2-His^6^, residues 1-116); full-length VAMP2^4X^ (VAMP2-His^6^, residues 1-116 with L70D, A74R, A81D, L84D mutation), full-length t-SNARE complex (mouse His^6^-SNAP25B, residues 1-206 and rat Syntaxin1A, residues 1-288); Synaptotagmin (rat Synaptotagmin1-His^6^, residues 57-421); and Complexin (human His^6^-Complexin 1, residues 1-134). All mutants including Syt1^F349A^ (F349A); Syt1^Q^ (R281A/E295A/Y338W/R398A/R399A); Syt1^DA^ (D309A, D363A, D365A) and SNARE^Q^ (SNAP25 K40A/D51A/E52A/E55A/D166A) were generated in the above described Syt1 and t-SNARE background respectively using the QuickChange mutagenesis kit (Agilent Technologies, Santa Clara, CA). All proteins were expressed and purified as described elsewhere (Mahal et al. 2002; Krishnakumar et al. 2013; Krishnakumar et al. 2011; Weber et al. 1998). Lipids, 1,2-dioleoyl-snglycero-3-phosphocholine (DOPC), 1,2-dioleoyl-sn-glycero-3-(phospho-L-serine) (DOPS), 1,2-dipalmitoyl-sn-glycero-3-phosphoethanolamine-N-(7-nitro-2-1,3-benzoxadiazol-4-yl) (NBD-DOPE), phosphatidylinositol 4, 5-bisphosphate (PIP2) were purchased from Avanti Polar Lipids (Alabaster, AL). ATTO647N-DOPE was purchased from ATTO-TEC, GmbH (Siegen, Germany) and lipophilic carbocyanine DiD (1,1’-Dioctadecyl-3,3,3’,3’-Tetramethylindodicarbocyanine Perchlorate) was purchased from Thermofisher Scientific (Waltham, MA). Calcium Green conjugated to a lipophilic 24-carbon alkyl chain (Calcium Green C24) was custom synthesized by Marker Gene Technologies (Eugene, OR)

### Liposome Preparation

t-SNAREs and VAMP2 (± Syt1) containing small unilamellar vesicles (SUV) were prepared using rapid detergent (1% Octylglucoside) dilution and dialysis method as described previously (Weber et al. 1998; Ji et al. 2010). VAMP2 (± Syt1) containing SUVs were subjected to additional purification on the discontinuous Nycodenz gradient. The lipid composition was 80 (mole)% DOPC, 15% DOPS, 3% PIP2 and 2% NBD-PE for t-SNARE SUV and 88% DOPC, 10% PS and 2% ATTO647-PE for VAMP2 (± Syt1) SUVs. To mimic physiological copy numbers of protein, we used an input of protein: lipid ratio as 1: 400 for t-SNARE, 1:100 for VAMP2 for physiological density, 1: 500 for VAMP2 at low copy number, and 1: 250 for Syt1. This was based on well-established parameters namely that the reconstitution efficiency for SNAREs and Syt1 is roughly 40-50% (densitometry analysis of the proteoliposomes) and only approximately 50–60% of the proteins are externally oriented (chymotrypsin protection analysis) (Ji et al. 2010; Ramakrishnan et al. 2019; Weber et al. 1998). Based on the densitometry analysis of Coomassie-stained SDS gels, we estimated vesicles at physiological density, contained 74 ± 4 and 26 ± 6 copies of outward-facing VAMP2 and Syt1 respectively (Figure 1 Supplement 1) and vesicles at low copy number of VAMP2 contained 13 ± 2 and 26 ± 6 copies of outward-facing VAMP2 and Syt1 respectively.

### Single Vesicle Fusion Assay

All the single-vesicle fusion measurements were carried out with suspended lipid bilayers as previously described (Ramakrishnan et al. 2019; Ramakrishnan et al. 2018). Briefly, t-SNARE-containing giant unilamellar vesicles as prepared using the osmotic shock protocol (Motta et al. 2015) were busted onto freshly plasma-cleaned Si/SiO2 chips containing 5 μm diameter holes in presence of HEPES buffer (25 mM HEPES, 140 mM KCl, 1mM DTT) supplemented with 5 mM MgCl_2_. The bilayers were extensively washed with HEPES buffer containing 1 mM MgCl_2_ and the fluidity of the t-SNARE containing bilayers was verified using fluorescence recovery after photo-bleaching using the NBD fluorescence. Vesicles (100 nM lipids) were added from the top using a pipette and allowed to interact with the bilayer for 10 minutes. We used the ATTO647-PE fluorescence to track vesicle docking, post-docking diffusion, docking-to-fusion delays and spontaneous fusion events. Subsequently, the excess vesicles in the chamber were removed by buffer exchange and 1 mM CaCl_2_ was added from the top to monitor the effect of Ca^2+^ on the docked vesicles. In all experiments, we used an inverted laser scanning confocal microscope (Leica-SP5) equipped with a multiwavelength argon laser including 488, diode lasers (532 and 641 nm), and a long-working distance 40X water immersion objective (NA 1.1). The emission light was spectrally separated and collected by photomultiplier tubes. To cover large areas of the planar bilayer and simultaneously record large ensembles of vesicles, the movies were acquired at a speed of 150 ms per frame t. Accurate quantification and fate of each vesicles were analyzed using our custom written MATLAB script.

### Single-Vesicle Docking Analysis

To get an accurate count of the docked vesicles, we used VAMP2 protein with mutations in the C-terminal half (L70D, A74R, A81D & L84D; VAMP2^4X^) that eliminates fusion without impeding the docking process. For the docking analysis, VAMP2^4X^ containing SUVs (vSUV^4X^) were introduced into the chamber and allowed to interact with the t-SNARE bilayer. After a 10 min incubation, the bilayer was thoroughly washed with running buffer (3x minimum) and the number of docked vesicles were counted. For an unbiased particle count, we employed a custom-written algorithm to count particles from top-left to bottom-right that ensures every spot is counted only once.

### Calcium Dynamics

To quantify the kinetics of Ca^2+^-triggered vesicle fusion, we used Ca^2+^-sensor dye, Calcium Green conjugated to a lipophilic 24-carbon alkyl chain (Calcium Green C24) introduced in the suspended bilayer to directly monitor of Ca^2+^ arrival. Calcium green is a high-affinity Ca^2+^-sensor (K_d_ of ∼75 nM) and exhibits a large fluorescent increase (at 532 nm) upon binding Ca^2+^. To accurately estimate the arrival of Ca^2+^ to/near the vesicles docked on the bilayer, we used confocal microscopy equipped with resonant scanner focused at or near the bilayer membrane and acquired movies at a speed of up to 36 msec per frame. We typically observed the fluorescence signal increase at the bilayer surface between about 3 frames (∼100 msec) after Ca^2+^ addition (Figure 2 Supplement 1). We therefore used 100 msec as the benchmark to accurately estimate the time-constants for the Ca^2+^-triggered fusion reaction.

## ACKNOWLEDGEMENTS

This work was supported by National Institute of Health (NIH) grant DK027044 to JER.

## AUTHOR CONTRIBUTIONS

SR, MB, JC performed the *in vitro* fusion experiments. SR, MB, JER, SSK designed the experiments, and analyzed the data. JER, SSK wrote the manuscript. All authors read and edited the manuscript.

---

**Figure 1 Supplement 1.**
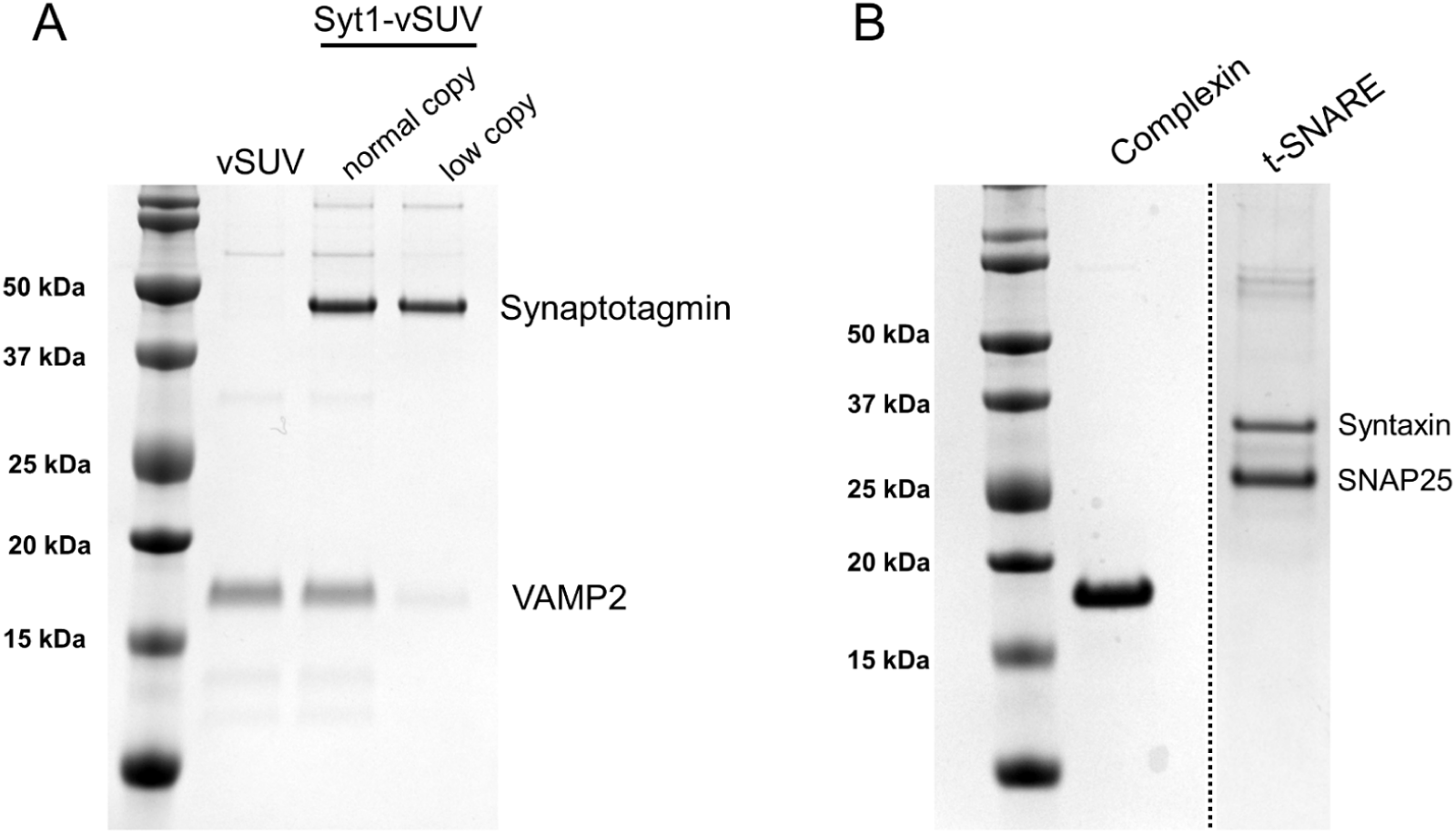
Coomassie-stained SDS-PAGE analysis of the proteins used in this study. (A) Gel image of the VAMP2 ± Syt1 vesicles. Based on the densitometry analysis, we estimated each vesicle contain 74 ± 4 and 26 ± 6 copies of outward-facing VAMP2 and Syt1 respectively under ‘normal’ physiologically-relevant condition and 13 ± 2 and 26 ± 6 copies of VAMP2 and Syt1 under low-copy conditions. (B) In all fusion experiments, co-purified t-SNAREs containing 1:1 complex of Syntaxin1a and SNAP25 was reconstituted onto the free-standing bilayer and full-length Complexin 1 was added in solution.

**Figure 1 Supplement 2.**
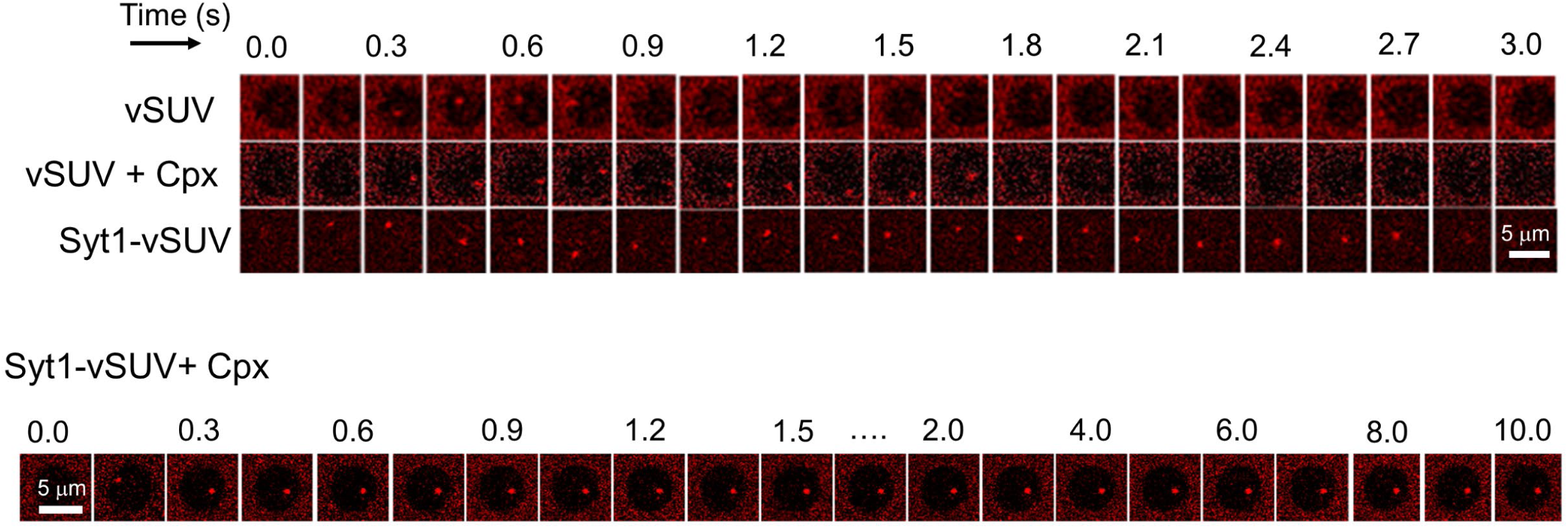
Representative time-lapse fluorescence (ATTO67N-PE) images of a docked SUVs showing that all VAMP2-containg SUVs (vSUVs) are diffusively mobile upon docking and fuse spontaneous within 1 sec. This behavior is un-altered upon addition of 2 μM Cpx (vSUV + Cpx). Inclusion of Syt1 in the v-SUVs (Syt1-vSUV) introduces a significant docking-to-fusion delay, but most vesicles proceed to fuse spontaneously. In contrast, Cpx, when added along with Syt1 (Syt1-vSUV + Cpx), produces a stable clamp and the vesicles are docked in-place and largely immobile and do not fuse during the observation period. Images corresponding to a single 5 μm suspended bilayer is shown.

**Figure 1 Supplement 3.**
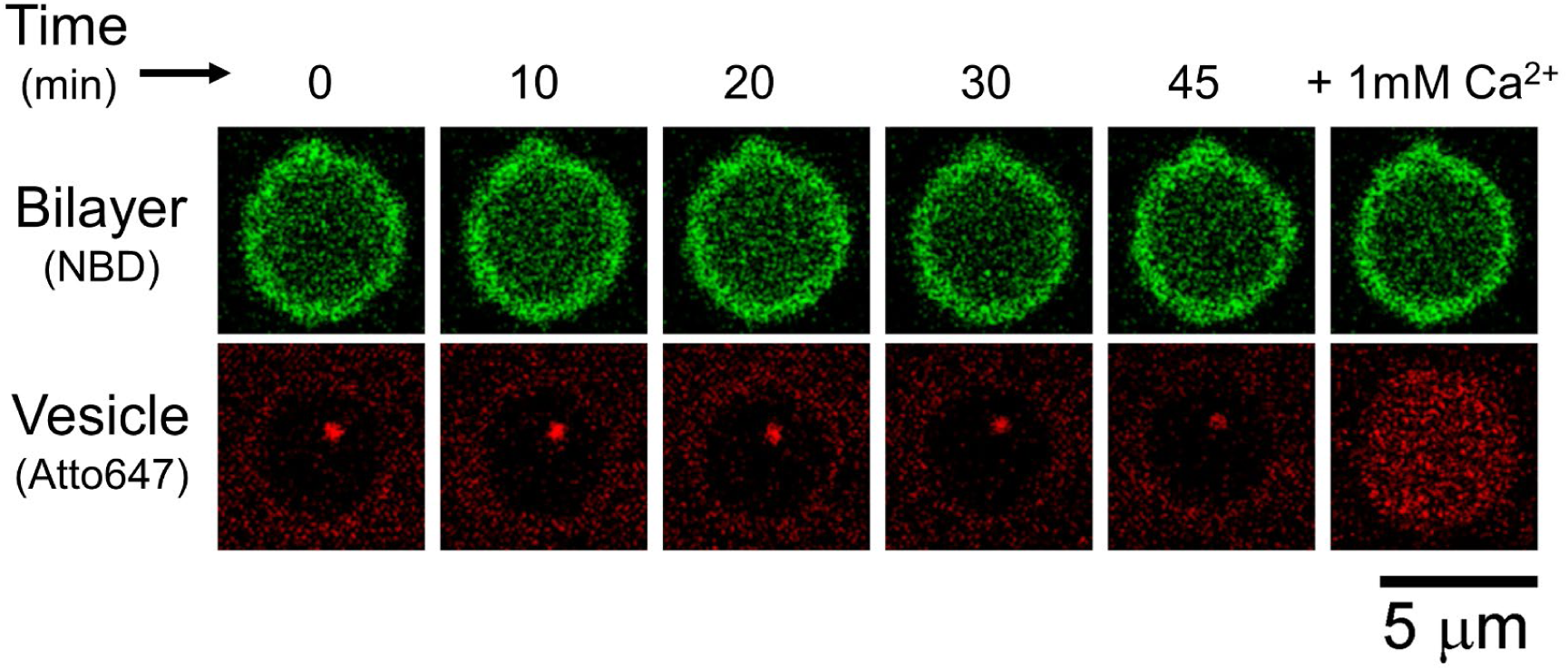
Representative fluorescence (ATTO647N-PE) image (bottom row) showing that the vesicles in the presence of Syt1 and Cpx dock and stay immobile at the site of attachment for up to 45 min and remain Ca^2+^-sensitive. To obtain these long time-lapse image series, the vesicles were imaged only every 10 min for up to 1h. At each time point, we also recorded the NBD-fluorescence (top row) to verify the stability of the bilayer. Images corresponding to a single 5 μm suspended bilayer is shown.

**Figure 2 Supplement 1.**
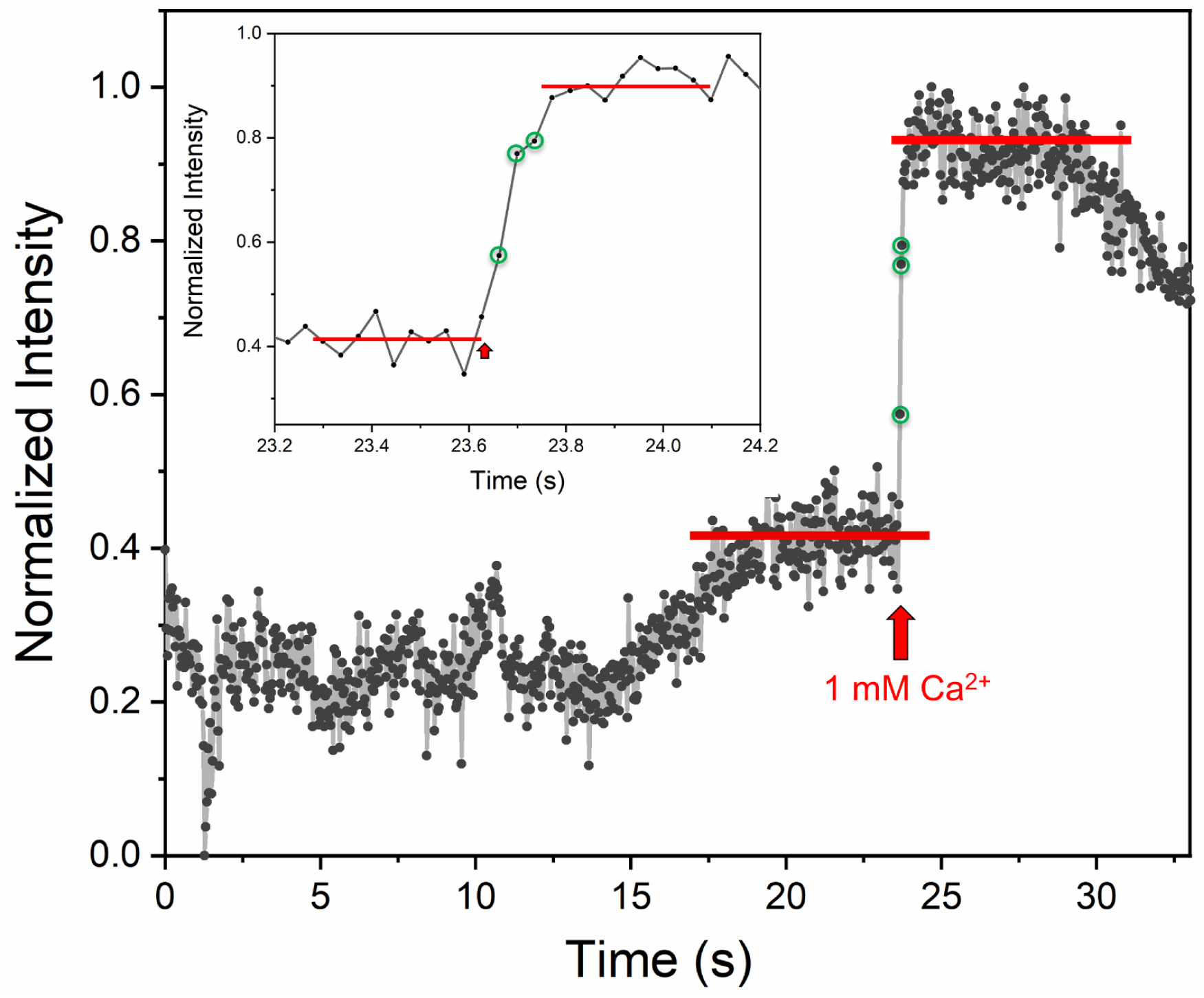
Ca^2+^-sensor dye, Calcium Green, introduced in the suspended bilayer (via a lipophilic 24-carbon alkyl chain) was used to monitor the arrival of Ca^2+^ at/near the docked vesicles. A representative fluorescence trace acquired at a speed of ∼36 msec per frame using confocal microscopy equipped with resonant scanner focused at or near the bilayer membrane is shown. The fluorescence intensity increase upon binding Ca^2+^ was observed ∼3 frames (∼100 ms) after the addition of Ca^2+^ at the top of chamber. This was used to benchmark and to accurately estimate the Ca^2+^-triggered fusion rate.

**Figure 2 Supplement 2.**
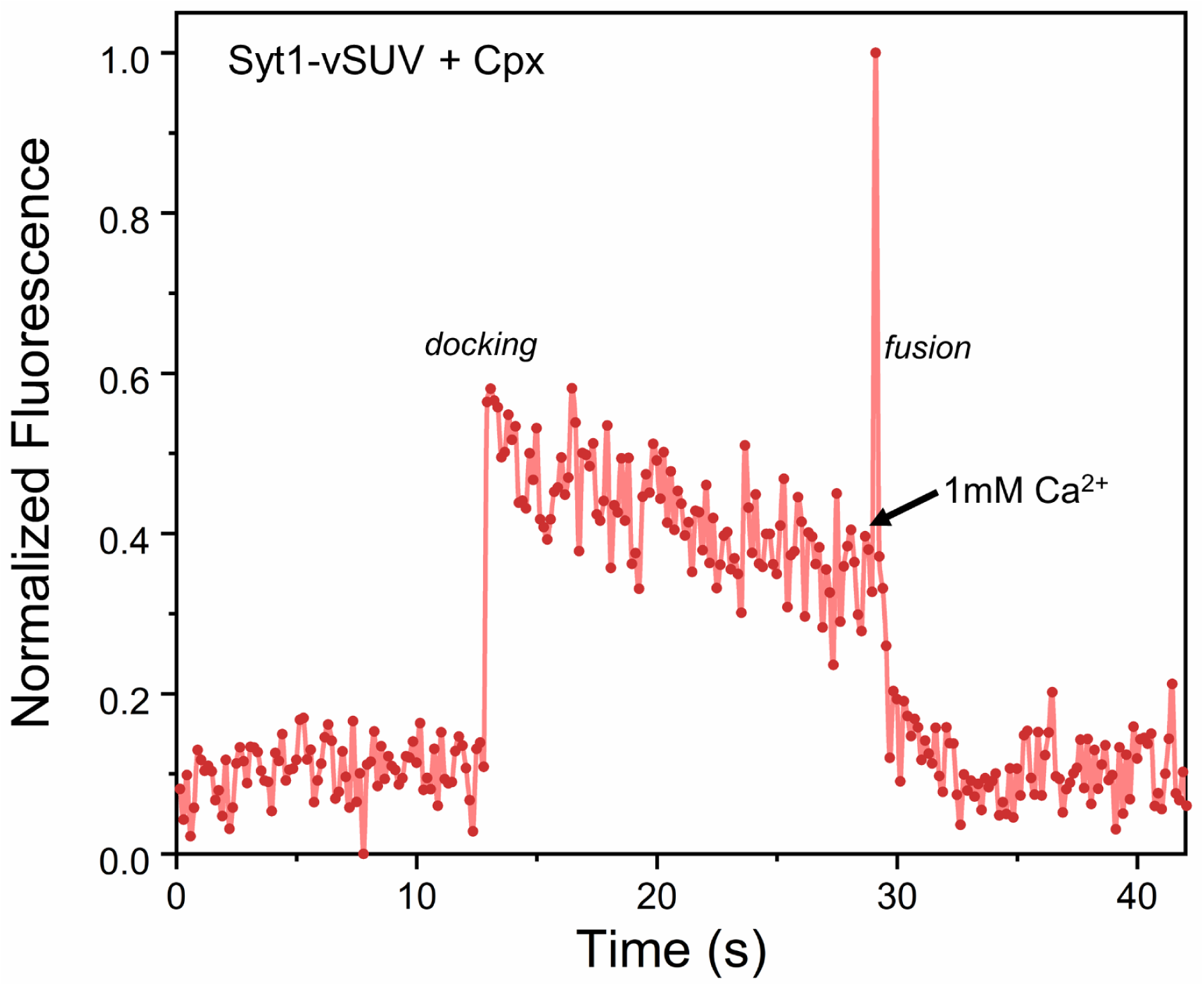
The effect of Syt1 and Cpx on SNARE-driven fusion assessed using a content-release assay with Sulforhodamine B loaded vesicle. Sulforhodamine B is largely self-quenched when encapsulated inside a SUV. Fusion of vesicle results in dilution of the probe, which is accompanied by increasing fluorescence. Representative fluorescence trace showing that in the presence of 2 μM Cpx, the Syt1-vSUV (loaded with Sulphorhodamine B) dock and remain in place, until triggered to fuse by Ca^2+^ (1mM) addition. This is in line with the lipid mixing data (Figure 2) showing that Syt1 and Cpx act together to establish Ca^2+^-regulated exocytosis.

**Figure 3 Supplement 1.**
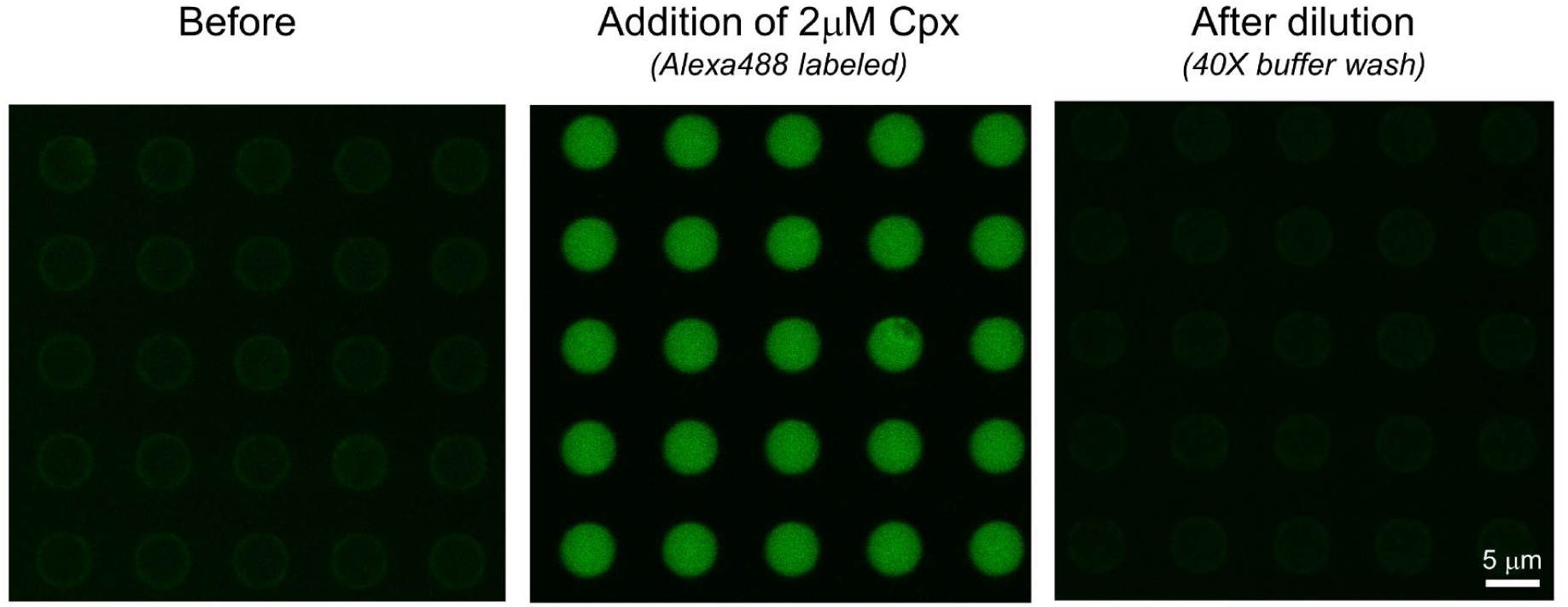
Representative fluorescence images showing that the extensive (40X) buffer wash results in complete washout of Cpx from clamped Syt1/Cpx vesicles. This was assessed using fluorescently (Alexa488) labeled Cpx and fluorescence recorded prior to the addition of Cpx (left panel), upon addition of Cpx (middle) and after dilution by 40X buffer wash (right panel) are shown. Images corresponding to multiple 5 μm suspended bilayer is shown. Note: The vesicle and bilayer fluorescence are not shown for clarity.

**Figure 3 Supplement 2.**
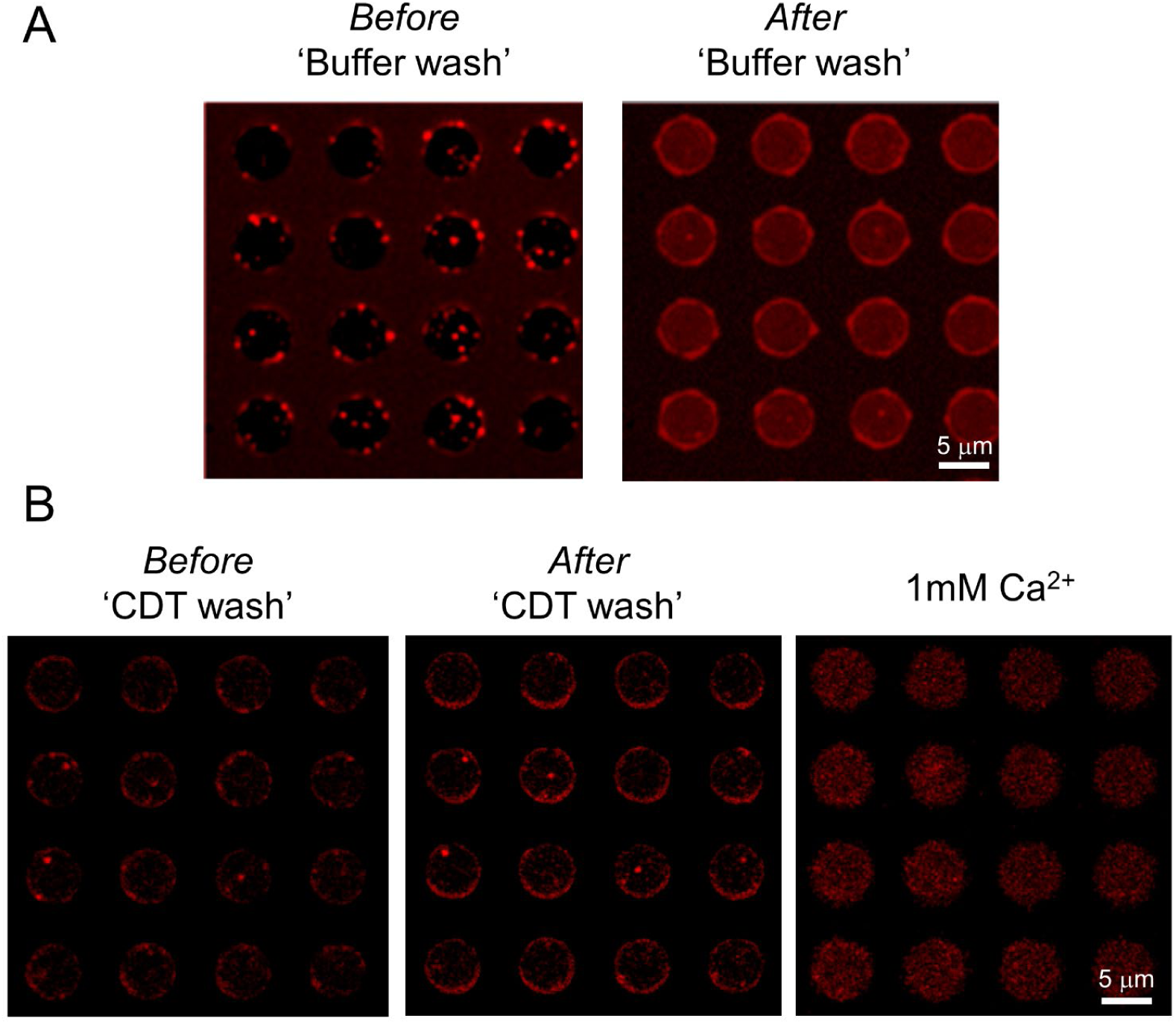
(A) Representative fluorescence (ATTO647N) image showing that clamped Syt1/Cpx vesicles (left panel) fuse spontaneously upon wash out of Cpx by dilution. Upon fusion, the ATTO647N dye on the vesicles mixes with suspended bilayer resulting in increase of the background ATTO647N fluorescence (right panel). (B) In the presence of excess inhibitory soluble cytoplasmic domain of the t-SNARE complex (CDT), the vesicles remain docked even after the removal of previously-bound Cpx by the buffer wash (middle panel). Subsequent addition of Ca^2+^ (1 mM) triggered rapid and synchronous fusion of these docked vesicles. Images corresponding to multiple 5 μm suspended bilayer is shown.

**Figure 4 Supplement 1.**
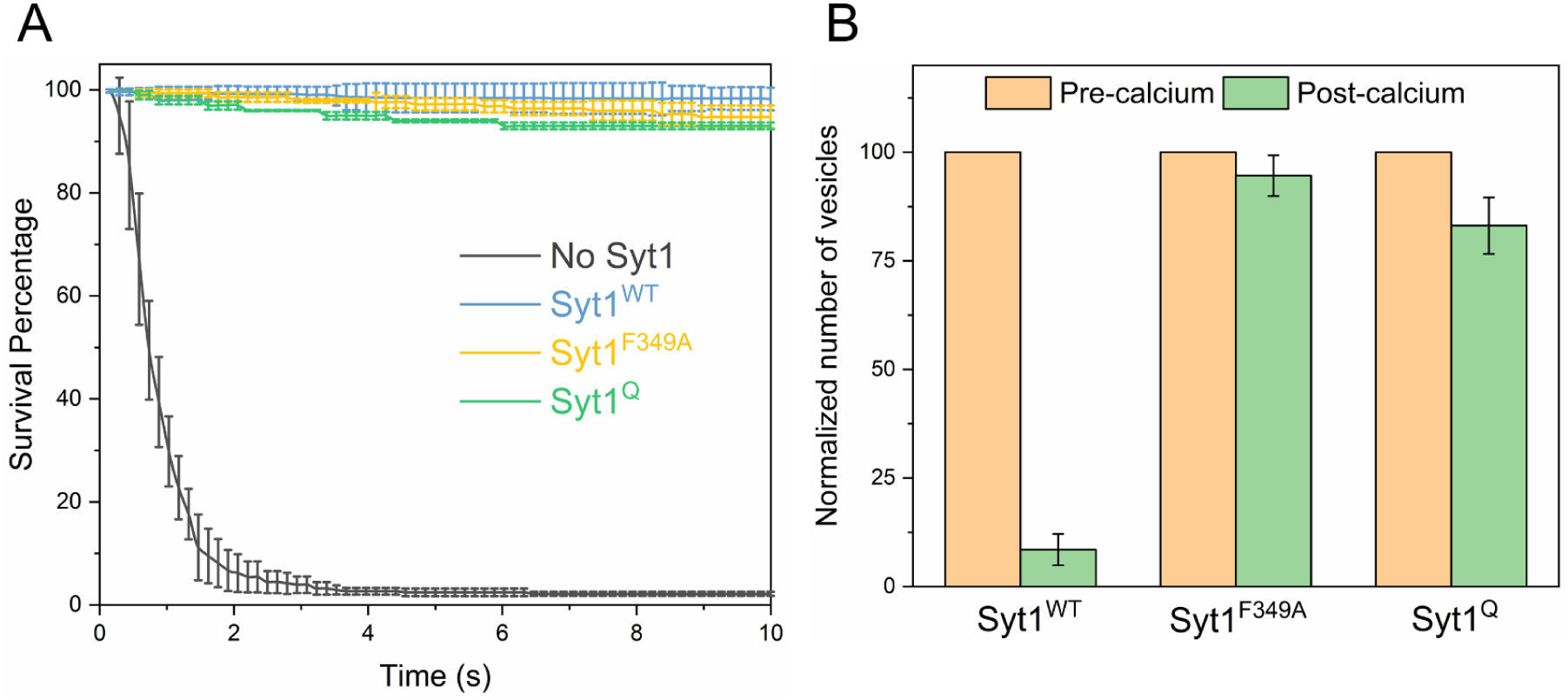
The effect of targeted Syt1 mutations were assessed under physiologically relevant conditions (∼74 copies of VAMP2 and ∼25 copies of Syt1) in the presence of 2 μM Cpx. (A) Survival analysis shows in contrast to the low copy conditions (Figure 4), disrupting the Syt1-SNARE primary interface (Syt1^Q^, green curve) or destabilizing Syt1 oligomerization (Syt1^F349A^, yellow curve) had no effect on fusion clamp in the presence of Cpx, with stably docked vesicles formed under all conditions. However, these vesicles are insensitive to Ca^2+^ and do not fuse upon addition of 1mM Ca^2+^ (B). This suggests that in the absence of the Syt1 clamp, Cpx blocks SNARE assembly irreversibly. The average values and standard deviations from 3 independent experiments are shown for each condition. In total, ∼200 vesicles were analyzed for each condition.

